# The Feasibility of Predicting Impending Malignant Ventricular Arrhythmias on the basis of Signal Complexity of Heartbeat Intervals

**DOI:** 10.1101/2019.12.25.888321

**Authors:** Zheng Chen, Koshiro Kido, Naoaki Ono, MD Altaf-Ul-Amin, Shigehiko Kanaya, Ming Huang

## Abstract

Malignant ventricular arrhythmias (MAs), such as ventricular tachycardia (VT) that presages cardiac arrest, present the highest hurdle for the healthcare community to overcome. Given that MAs occur unpredictably and lead to emergencies, convenient tracking devices, e.g. photoplethysmogram (PPG), that could predict MAs would be irreplaceably valuable. Since the use of heartbeat intervals (HbI) to predict the occurrence of arrhythmias is becoming more feasible, a further attempt to establish a new convenient approach for predicting impending MAs with HbI is worth trying. Assuming that intrinsic characteristics of MAs (VT and ventricular fibrillation: VF) can be revealed by a suitable approach on the basis of signal complexity, we propose an approach that first expresses the physiological status of the heart by HbI; then delineates the patterns of HbI by a new complexity metric (refined composite multi-scale entropy: RCMsEn); and finally trains a nonlinear machine learning model (random forest: RF) to learn the specific patterns of MAs so as to differentiate them from the normal sinus heart rhythm(N) and other prevalent arrhythmias (atrial fibrillation: AF, and premature ventricular contraction: V).

For calculating entropy values and predicting MAs as early as possible (which is the aim of this study), two specifications are of interest: the minimal length of HbI needed to delineate the MAs patterns sufficiently (*len_min_*), and the maximum time length at which our model can predict impending MAs (*time_max_*). We compared the RF model with support vector machine (SVM) models based on linear and Gaussian kernels. Results show that the RF model performs the best, reaching a 99.24% recall and a 99.87% precision for a HbI of 500 heartbeats (the *len_min_*) 374 seconds (the *time_max_*) preceding the occurrence of MAs. The HbI samples in this study were extracted from an electrocardiograph (ECG). However, given the subtle difference (0.1 ms typically) between the R-R interval of ECG and the P-P interval of PPG, this approach could be extended to HbI acquired by the PPG sensor and thus should be of substantial theoretical and practical significance in cardiac arrest prevention.

## I. INTRODUCTION

THE cardiovascular system is controlled by complex self-regulation [1], involving various physiological parameters such as blood pressure and body temperature. The nonlinear regulation of the heart may be attributed to the complexity of the fractal-like structure of the His-Purkinje fiber network [2]. Defects in the afferent system or electrical conduction system may plausibly give rise to behavioral difference that can be manifested by signal complexity. Moreover, the cellular defects of the cardiomyocyte that cause cardiac arrhythmias can also be manifested by signal complexity changing [3].

Metrics defined on the basis of entropy are appropriate to qualify the complexity of signals, and a series of modifications and validations for entropy-based metrics has been made. Sample entropy (SampEn), which is a modification over the approximate entropy by excluding the self-match, has been used to analyze heart rate variability [4]. However, SampEn requires samples of 10^*m*^ ~ 30^*m*^ in length (*m*: the length of a compared run of data) and lacks consistency in some cases. To solve the issues of SampEn, Costa *et al*. proposed multi-scale entropy (MsEn), in which the original signal is coarse-grained by moving the average over different scales without overlapping and the entropy values are calculated for each scale [5], [6]. MsEn enables a deeper investigation of the signal of interest with multi-resolution and generates more stable results [7]. However, this study requires a relatively long signal. More recently, the refined composite multi-scale entropy (RCMsEn) was proposed to relax the requirement of signal length [8], [9].

It has become clearer that many kinds of cardiac arrhythmias can be characterized by nonlinear dynamics [10], so researchers have been trying to characterize arrhythmias of different origins so as to distinguish them by nonlinear metrics of the heartbeat interval (HbI). The prospect of this has become clearer with the progress in wearable devices and the Internet of Things (IoT). Owis *et al*. succeeded in distinguishing normal sinus heart rhythm from abnormal rhythm by using the correlation dimension and Lyapunov exponents [11]. Zhou *et al*. tried to use the Shannon entropy in detecting the atrial fibrillation from normal sinus heart rhythm and showed promising results in a real-time application [12]. cite The HbI is canonically extracted from the R-R interval of an electrocardiograph (ECG), which cannot be acquired easily outside a hospital. Fortunately, HbI has been validated to also be able to be obtained by another sensing technology, e.g., the peak-to-peak interval of photoplethysmography (PPG) [13]. The diversification of the signal source endows the methods based on HbI a much broader field of applications, from the clinical setting to personal healthcare [14]. This is because the wearable/unconstrained measurements provide more flexibility and more dynamic information at the expense of a lower signal quality due to improper sensor settings, body movement, etc. [15]–[17]. Sometimes, the signal is so severely contaminated by noise that only the heart rate information (e.g., from the R-R interval of ECG or P-P interval of PPG) can be extracted accurately. Hence, a reliable approach using the metrics derived from HbI solely to reflect the physiological/pathological status of the heart is a practical necessity [18], [19].

This necessity should be especially strengthened in detecting malignant ventricular arrhythmias (MAs), which are life-threatening and may develop without specific cardiac disorders, e.g., long QT syndrome and Wolff-Parkinson-White (WPW) syndrome. An efficient approach, from acquiring physiological signals to decision-making using the signal, to predict impending MAs will be incomparably valuable for the healthcare community [20], [21].

Among the different types of MAs, the most common type is ventricular tachycardia (VT), which may presage cardiac arrest, and the most lethal type, meaning a preterminal event, is ventricular fibrillation (VF). Therefore, in the past two decades, we have seen a number of studies that have tried to predict VT and VF using the statistical characteristics of heart rate [22], frequency-domain analysis of heart rate variability (HRV) [23], and the combination of HRV metrics and artificial neural networks (ANNs) [24]. More recently, Lee *et al*. constructed an ANN binary model using 14 parameters from HRV and respiratory rate variability to predict VT occurrence and claimed that their model can predict VT 30 seconds prior to the event with a sensitivity of 0.88, a specificity of 0.82, and an area under the receiver operating characteristic curve (AUC ROC) of 0.93 [25]. Taye *et al*. constructed an ANN model with morphological features of ECG signal to predict VT 30 seconds prior to its onset and claimed that the accuracy of their approach reached 98.6% [26].

Although these studies demonstrate the possibility of MA prediction, we need to recognize that some critical issues remain. The first is the signal source. Regarding the contingency of MAs, approaches that demand high quality ECG signals will not be the best choice. Second, in considering the similarities of arrhythmias in a feature space, a binary model that separates the MAs from normal sinus rhythm is not sufficient. Considering all these achievements and remaining issues in MA prediction, our assumption and the aim of this study are as follows:

Assumption: Autonomic regularization takes different measures to compensate for arrhythmias originating from different chambers of the heart. At a certain time prior to MAs occurring, the heart tries to take measure to compensate, which will result in specific activity patterns of the heart that are embedded in HbI. By using proper metrics, these specific activity patterns of MAs can be manifested and are unique enough to be separated from the activity patterns of normal sinus rhythm (N), and other common arrhythmias (atrial fibrillation: AF and ectopic ventricular heartbeat: V).

Aim: By manifesting the unique patterns of the HbI prior to the VT/VF event (preVT/preVF) with proper metrics of signal complexity, we aim to separate the preVT and preVF by a proper machine learning model from other heart rhythms. That is, we aim to construct a supervised machine-learning model for a multi-class classification problem. Especially, we focused on two specifications of our approach: the minimal length of HbI needed to delineate the MAs patterns sufficiently (*len_min_*), and the maximum time length at which our model can predict impending MAs (*time_max_*).

## II. MATERIALS AND METHODS

The concept of our approach was implemented in three consecutive steps, which will be introduced in II-A~C. In II-A, we introduce the acquisition of signal samples and the necessary pre-processing. Since the VT/VF-related samples are difficult to acquire by measurement, we used a well-established public dataset, the Spontaneous Ventricular Tachyarrhythmia Database^1^ (SVTDB, Version 1.0 from Medtronic Inc), to validate our assumption. II-B gives the details of the nonlinear metrics that we used to manifest the activity patterns of different heart rhythms, as well as the details about feature calculation. Finally, in II-C we chose the best model by comparing the performances of support vector machine (SVM) models of linear and Gaussian kernels and a random forest (RF) model in using the nonlinear metrics for preVT/preVF classification.

### A. Data and preprocessing

The preVT and preVF samples, which were recorded by implanted cardioverter defibrillators (ICD), were extracted from the SVTDB [27]. The SVTDB contains 135 pairs (a spontaneous episode of VT or VF and an intrinsic normal sinus rhythm episode) of R-R interval (RRI) time series, which were recorded in 78 subjects. Since the ICD (Medtronic Jewel Plus TM ICD 7218) possesses a buffer containing the 1024 most recently measured RRIs, the 1024 RRIs that immediately precede the detected event (VT or VF) could be captured. Corresponding to the type of MA events that the heart developed, the 1024 RRIs are defined as preVT or preVF. Noteworthily, the HbI is represented by RRI here. The prediction of MAs is reflected in the preVT/preVF selection, whose scheme is reflected in Fig. 1. Specifically, the foremost RRIs (length: 300~1000) were extracted in such a way that the series was taken out from the head as the window slid down with a stride equaling 1 RRI, and the sampling stopped when 25 samples were taken out. This scheme resulted in a prediction prior to the MA event at a time length from 700 RRI (when the length of RRI is 300) to 1 RRI (when the length of RRI is 1000), about 525~1 seconds under 80 bpm.

**Fig. 1.**
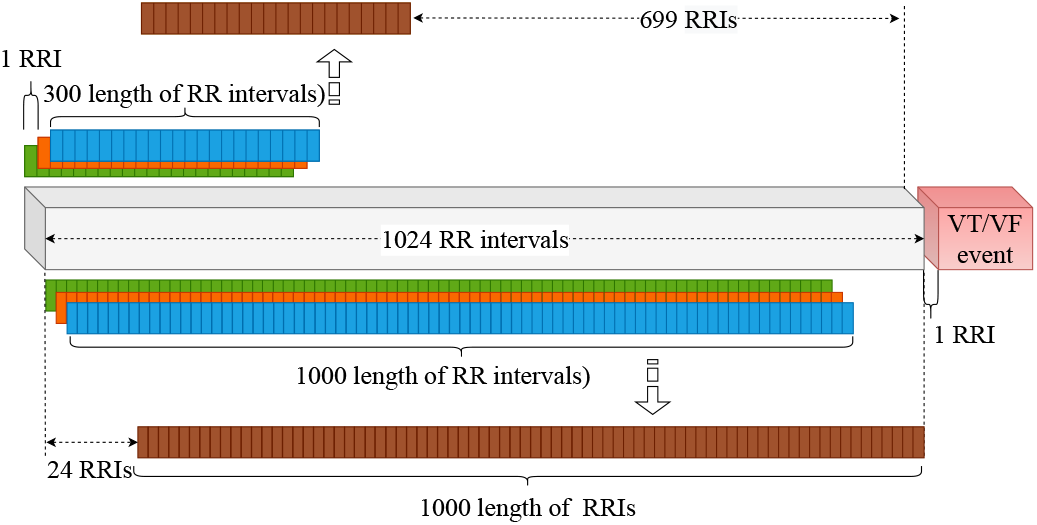
Illustration of extraction of preVT and preVF samples.The samples were extracted at the foremost part of the 1024 RRIs, and the windows with a fixed length slid down with a stride of 1 until 25 samples were taken out.

To validate out assumption that the preVT and PreVF have their own unique activity patterns that are distinguishable from other rhythms, we extracted normal sinus heart rhythm (N) and two prevalent types of arrhythmias (atrial fibrillation: AF, and ventricular ectopic heartbeat: V) from the MIT-BIH arrhythmia database (MITDB)^2^. Since each record in this database has been annotated by two physiologists for each heartbeat in view of the ECG morphology of the waveform, and the annotations have been adjusted to the R-wave peaks, the RRI samples were extracted on the basis of the R-peak annotations [28]. By using the manual annotations of the heartbeats in the database, samples of each rhythm are extracted as follow:

- samples of N type contain normal heartbeat rhythm throughout the sample only;
- samples of AF type contain at least 10% (*σ_AF_*) atrial fibrillation beats;
- samples of V type contain at least 5% (*σ_V_*) ventricular ectopic beats.

### B. The nonlinear metrics (RCMsEn) and sample calculation

The RCMsEn was used to calculate the nonlinear metrics that reflect the complexity of a HbI signal because it provides multi-resolution information and works well for a relatively short signal. The relevant SampEn and MsEn will be introduced briefly to facilitate a quick grasp of RCMsEn.

Suppose there is a short signal *X* = (*x*_1_, *x*_2_, …, *x_n_*). The SampEn is defined by

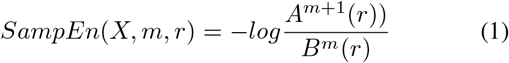

where *m* is the embedding dimension, which is set to 2 in most of the cases; the *r* is the tolerance and is usually set as 0.1 0.2 of the standard deviation of the signal. In accordance with the setting of *m*, the signal is divided into fragments of a length of m and m + 1. The *A*^*m*+1^(*r*) is the number of pairs[*X̂*_*m*+1_(*i*),*X̂_m+1_*(*j*)]((*i* ≠ *j*, *X̂_m+1_*(*i*) : the ith fragment of *m* + 1 length), whose distance is shorter than *r*.

The MsEn was proposed to provide multi-resolution information on the basis of the entropy theory, in which the original signal is averaged over different scales defined by *τ* (scale *kϵ*[1; *τ*]). Therefore, for each scale *k*, a new time series is generated as

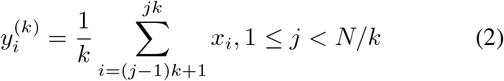

For each scale, the generated time series 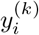 is used to calculate the SampEn. Hence,

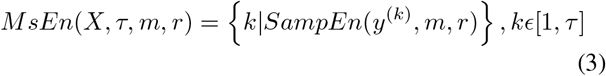

The entropy values of each scale together give us the chance to inspect the self-similarity of the signal from different time scales. However, the coarse-graining of the MsEn would decrease the length of the signal generated by a ratio of the scale.

RCMsEn is a modification based on the MsEn to solve the issue of data-length for the short signal, which generates k new time series 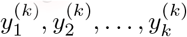 for the corresponding scale k by shifting the coarse-graining procedure to the right as shown in Fig. 2.

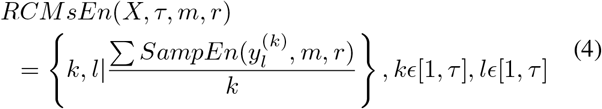

where the SampEn values are calculated individually and averaged over *k* for each scale. RCMsEn has been validated to provide more stable and consistent values, for especially short time series [9].

**Fig. 2.**
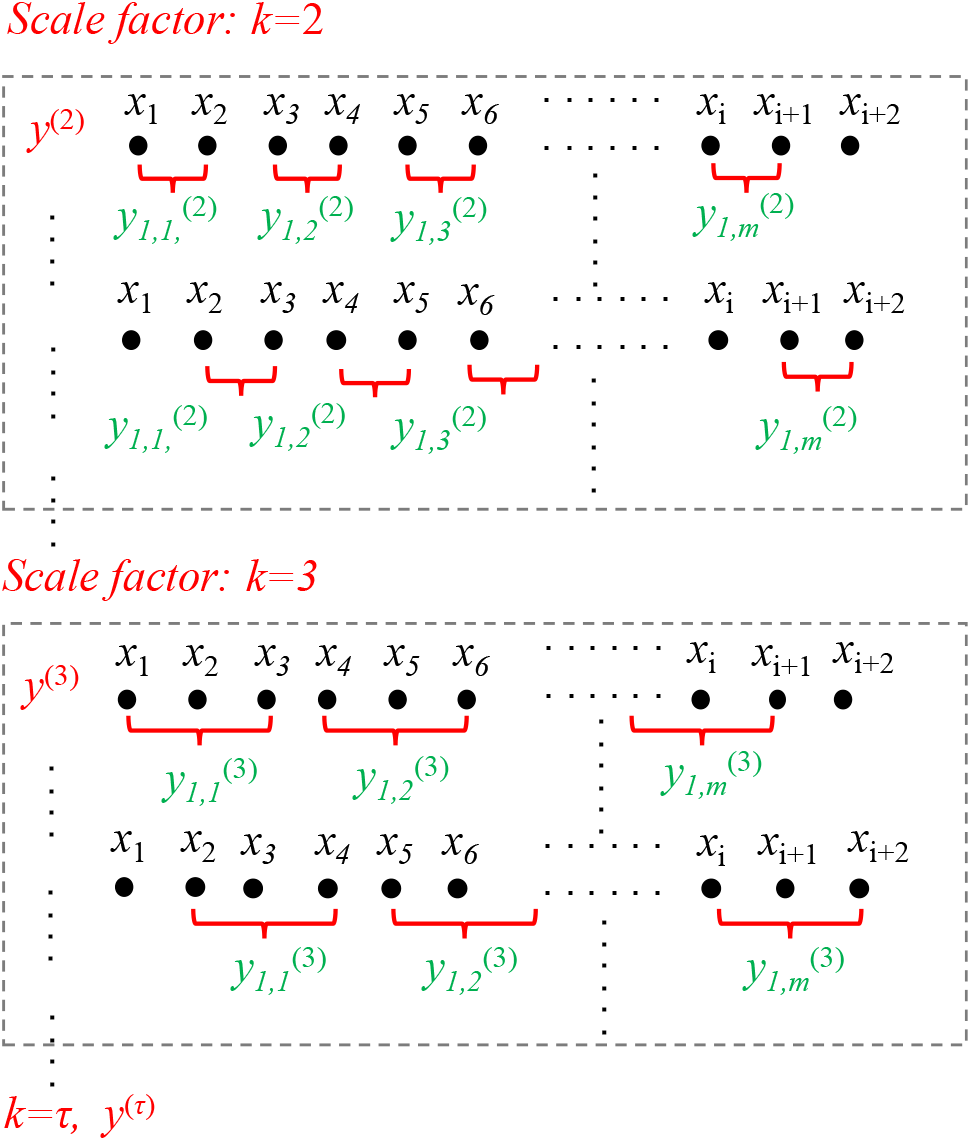
Illustration of calculation of RCMsEn.The process shown in the broken-line box repeats until the pre-defined is reached.

As mentioned above, the minimum length of the HbI signal (*len_min_*) and the corresponding maximum time length at which the model can predict impending MAs (*time_max_*) are of interest in this study. Therefore, we generated the RCMsEn values for HbI signals of different data-lengths: 300~1000 RRIs with 100 intervals.

The maximum scale *τ* of RCMsEn was set as 20. However, the available dimensions of the relatively short HbI, e.g. 300, may be fewer due to the undefined entropy. Therefore, as the input into the machine-learning model, the dimensions of the RCMsEn vectors of all the five types of heartbeats were aligned to the minimum dimensions. Noteworthily, even for the same type of heartbeat, the dimension may vary over samples, so the maximal scale was chosen in a way that if more than 1% of the samples showed undefined entropy, the scale was excluded.

### C. Machine-learning models

Models based on SVM (linear and Gaussian kernels) and RF were constructed separately to select the best model. A SVM is a discriminative classifier formally defined by a separating hyperplane. In other words, given labeled training data (supervised learning), a SVM outputs an optimal hyperplane that categorizes new examples. In two-dimensional space, this hyperplane is a line dividing a plane in two parts. A kernel is a way to place a feature space into a higher dimensional space to facilitate the separation of data. The kernel can have a linear or nonlinear form, whereas the dimension of the space can be finite (linear, Polynomial) or even infinite (Radial Basis Function: RBF). The RBF kernel can be described by

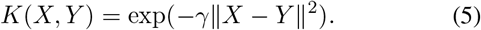

The RBF kernel is termed a Gaussian kernel when 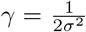, where the *σ* is a free parameter.

The RF is an ensemble learning algorithm that grows a number of tree-based weak classifiers to prevent the overfitting problem that can often be seen in a single complicated model. At the same time, the RF reduces the predictive variance by decorrelating the individual weak classifier by randomly selecting partial independent variables to grow a tree and by growing it with different bootstrapped samples. The RF was implemented as follows:

Construction of the RF ensemble

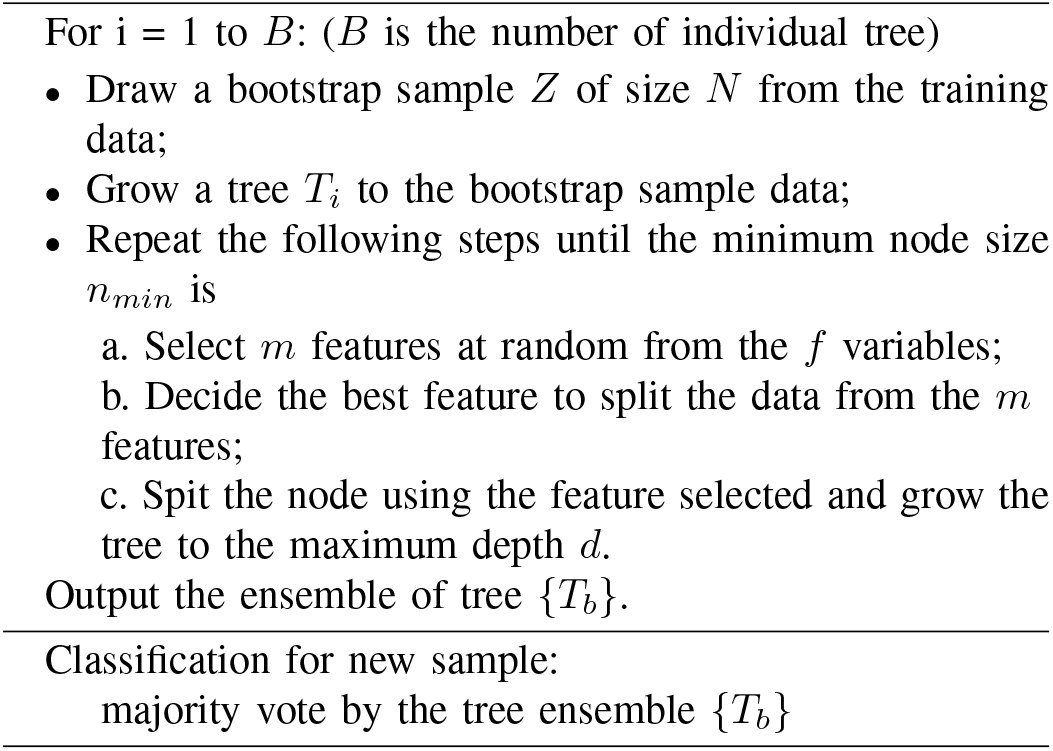

The RF reduces the variance of the ensemble in accordance with the following equation:

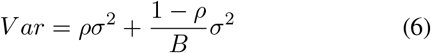

where the *ρ* is the correlation of trees and *σ*^2^ is the variance of the features (assumed as a constant for all features). By increasing the tree numbers, the second term on the right-hand side becomes minor; whereas the first term can be decreased by reducing the correlation of trees by a random selection of the m features.

The hyperparameters for the RF are the number of trees, max features, and max depth of a tree. If the computation time is not a parameter of concern, the larger the tree number (500 in this study), the smaller the variance. As mentioned above, only a portion of the independent variables (features) is used to grow a tree, and the number is defined by the maximum features 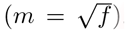. Finally, a tree with a deep depth may cause the overfitting problem, so the max depth of a tree must be defined. (*d* = 10 in this study).

The SVM and RF were chosen to generate the machine-learning models to confirm our observation of the analysis of RCMsEn using principle component analysis (results not shown in this paper). No clear separation can be seen for the 5 types in the first 3 principal components, and the entropy values of each scale that belong to the same type do not follow a normal distribution. The difficulty in applying linear methods may be due to the high variances in some features. These observations give us a hint that the RF model may outperform SVM models and that this can be confirmed by the result.

### D. Evaluation

The flow of this study from the HbI extraction to models construction and comparison is shown in Fig. 3. The workflow introduced above conceptually splits the feature space consisting of the RCMsEn values into the five types of heart rhythms and can be instantiated by machine learning models. To train and test the models, training samples and test samples were split at a ratio of 8:2 for each type of heartbeat. To evaluate the combined performance of the RCMsEn-based features and the classification models, class-specific precision and recall and specificity are used.

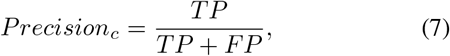

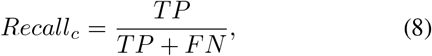

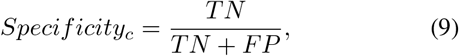

where subscript *c* denotes the class-wise calculation.

**Fig. 3.**
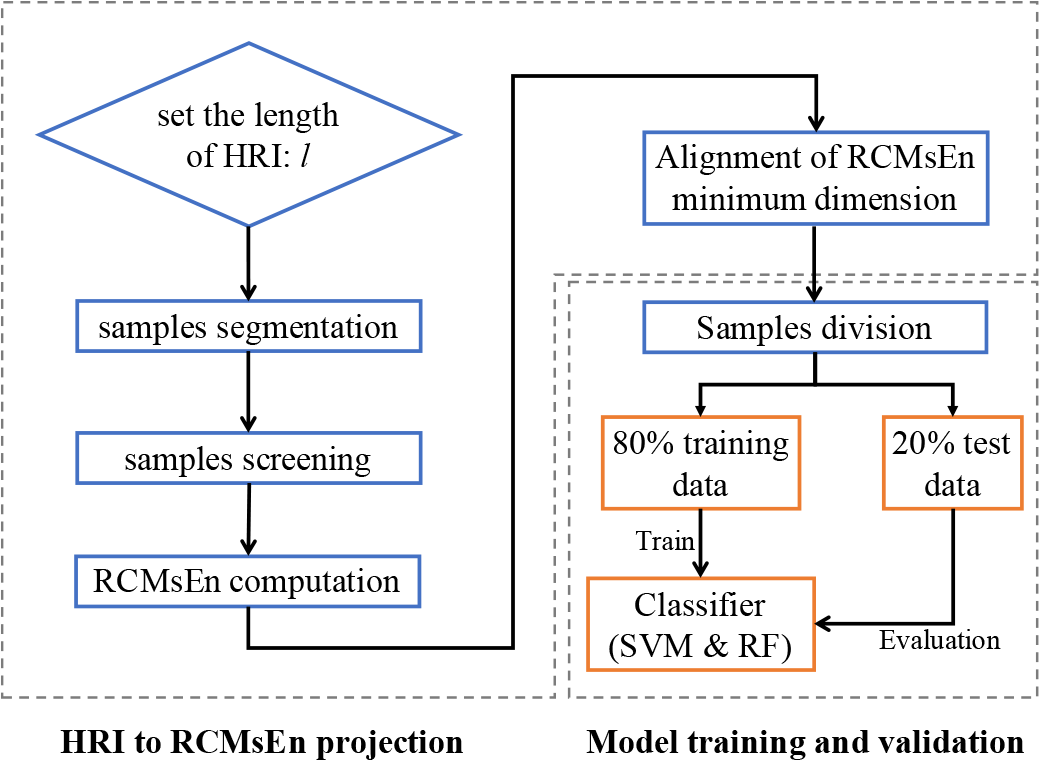
Main flow of the approach proposed in this study.The 20% test data are completely unseen in the training section.

A deeper understanding of the features of different scales on the classification can be revealed by analyzing the importance of the RF model. For each length of HbI, the importance of each scale can be obtained from the RF model. For the first nine important scales in terms of median, the Mann-Whitney *U* Test is used to test the significant difference between two scales in view of the small number of samples.

## III. RESULT

### A. Samples and Features based on RCMsEn

The undefined entropy still exists in samples of relatively short length. The scales available for HbI of different lengths are tabulated in TABLE I.

**TABLE I.** AVAILABLE SCALES OF RCMSEN FOR HBI OF DIFFERENT LENGTHS

| Length of HbI | Available scales |
| --- | --- |
| 300 | 1~3 |
| 400 | 1~6 |
| 500 | 1~11 |
| 600~1000 | 1~20 |

The overall trends of the five types are shown by their mean values in Fig. 4, which shows that preVT and preVF have similar trends but different trends from the other three. Moreover, all the five types show very consistent trends in the three different lengths. We have to point out that, although the five types show clear trends in terms of mean values, they are still difficult to classify due to the large variances. Furthermore, the ventricular arrhythmias intriguingly have lower entropy values on most of the scales than atrial and normal ones.

**Fig. 4.**
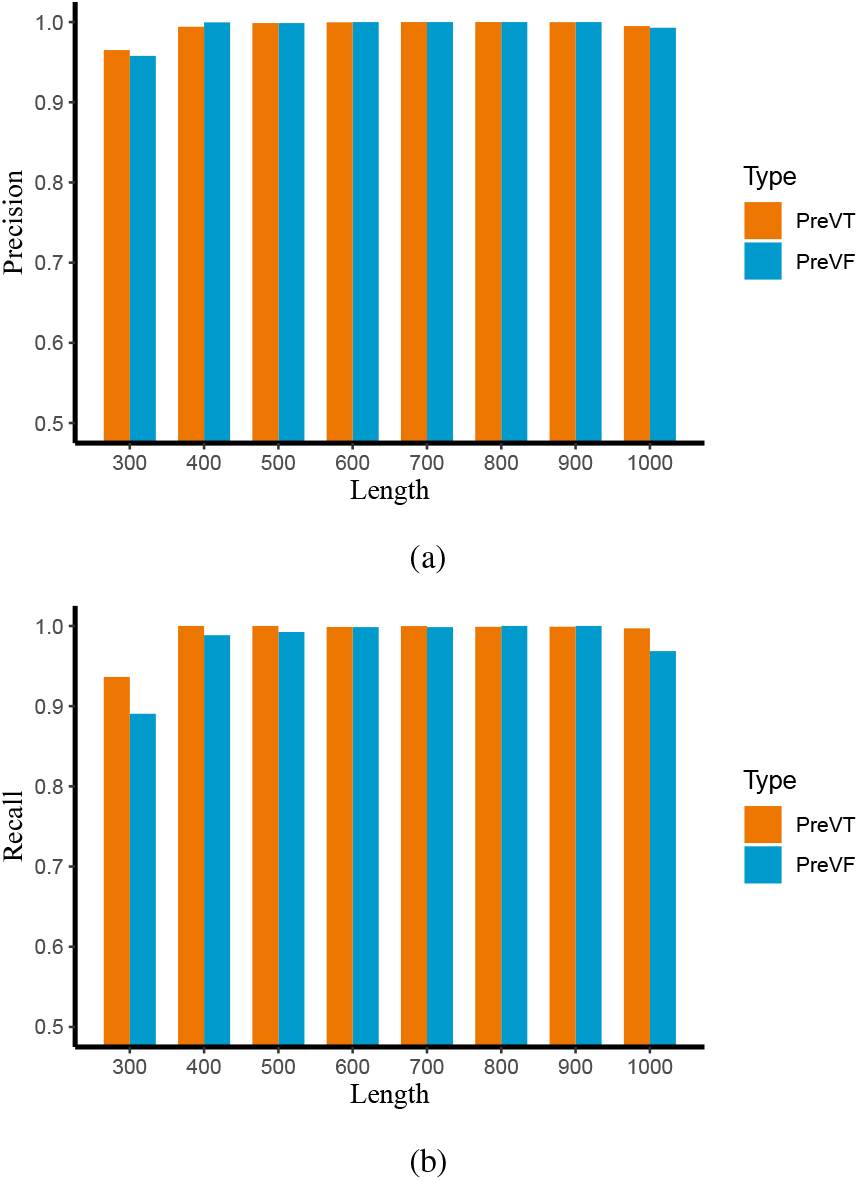
Precision (a) and Recall (b) for preVT and preVF detection.The values change significantly with the length of the HbI in the relatively short signals (300~500).

**Fig. 5.**
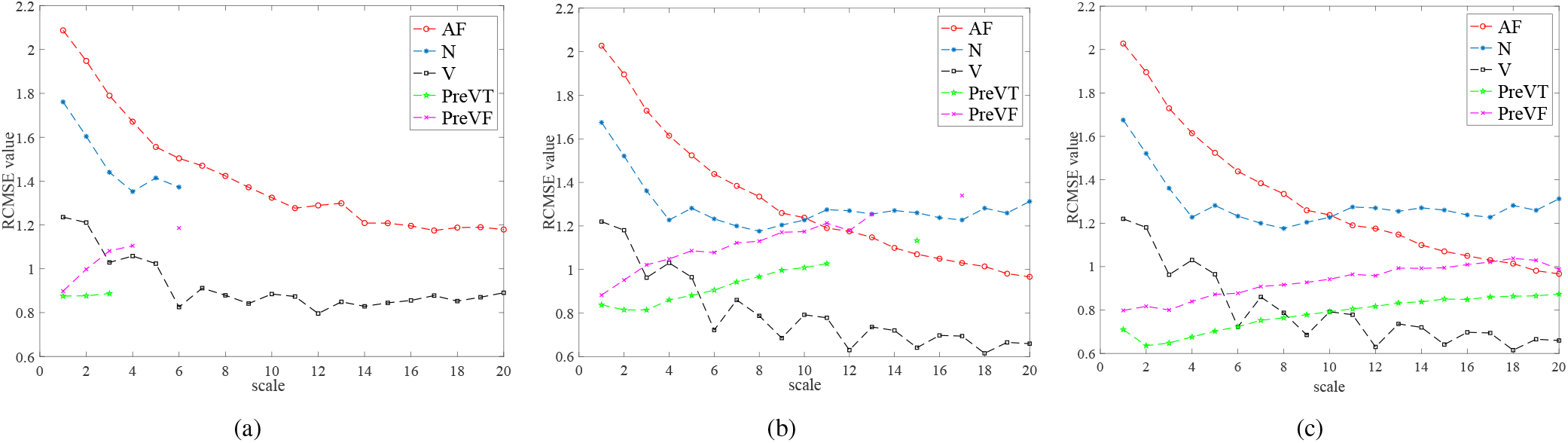
Means of RCMsEn for (a) 300, (b) 500, and (c) 1000 HbI.

### B. Model comparison for different lengths

The dimension of the RCMsEn features increases with the data-length as shown in TABLE I and is beneficial for the models to improve their performances. The effect of the dimension of features can be seen in TABLE II, which shows the performances of the SVM models and RF model in detail. The RF model using 500 RRIs performs the best; whereas the SVM models perform much worse, especially for preVF detection. Judging from the overall performances, we conclude that the RF model is the most appropriate classifier for this study.

**TABLE II.**
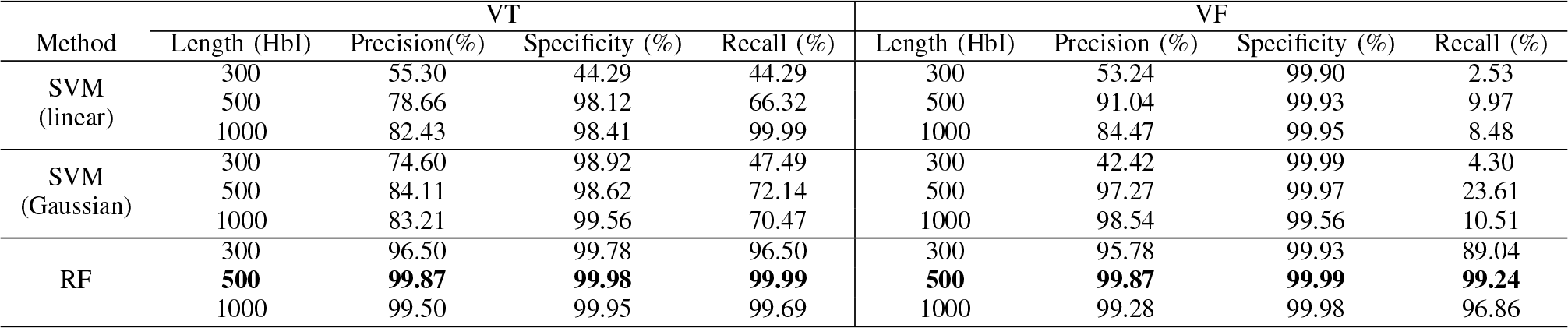
PERFORMANCES OF SVM MODELS AND RF MODEL

| Method | VT |  |  |  | VF |  |  |  |
| --- | --- | --- | --- | --- | --- | --- | --- | --- |
|  | Length (HbI) | Precision(%) | Specificity (%) | Recall (%) | Length (HbI) | Precision (%) | Specificity (%) | Recall (%) |
| SVM<br>(linear) | 300 | 55.30 | 44.29 | 44.29 | 300 | 53.24 | 99.90 | 2.53 |
|  | 500 | 78.66 | 98.12 | 66.32 | 500 | 91.04 | 99.93 | 9.97 |
|  | 1000 | 82.43 | 98.41 | 99.99 | 1000 | 84.47 | 99.95 | 8.48 |
| SVM<br>(Gaussian) | 300 | 74.60 | 98.92 | 47.49 | 300 | 42.42 | 99.99 | 4.30 |
|  | 500 | 84.11 | 98.62 | 72.14 | 500 | 97.27 | 99.97 | 23.61 |
|  | 1000 | 83.21 | 99.56 | 70.47 | 1000 | 98.54 | 99.56 | 10.51 |
| RF | 300 | 96.50 | 99.78 | 96.50 | 300 | 95.78 | 99.93 | 89.04 |
|  | **500** | **99.87** | **99.98** | **99.99** | **500** | **99.87** | **99.99** | **99.24** |
|  | 1000 | 99.50 | 99.95 | 99.69 | 1000 | 99.28 | 99.98 | 96.86 |

Focusing on the RF model, we further investigated the effect of the data length on performance. According to TABLE II, there are only very subtle differences in the range from 500 to 900 HbI; whereas 1000 HbI has somewhat inferior results. In view of the scale of 500 HbI going up to 11, the results suggest that the lower scales have more important features than the higher ones. Moreover, although the performances are stable from 500 to 900 HbI, 500 HbI is chosen because it corresponds to the earliest MA detection: *time_max_* = 374 seconds.

The confusion matrix in Fig. 6 gives more details about the RF model with 500 HbI. It can be seen that N, AV, and V types are clearly separated without mismatching. Focusing on the ventricular ones, the model is still able to provide an accurate stratification that separates the MAs from the relatively benign arrhythmias. Even inside MAs, the subclassification is accurate, and only one misclassification occurred.

**Fig. 6.**
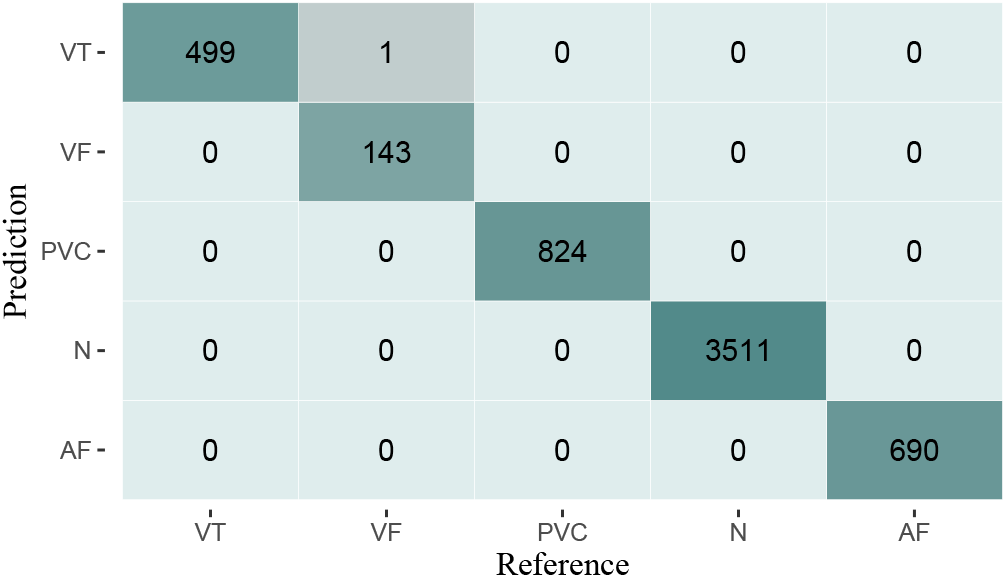
Confusion matrix of RF model input with 500 HbI. N, AF, and V are separated accurately without misclassification, and only one VT is mistakenly classified as VF.

### C. Importance of the features

It can be inferred from the results of the RF model using HbI of a length ≥ 500 that the finer scales (1~11) are important features for classification. The medians of normalized importance of the features are plotted in Fig. 7, which shows that the lower scales (*τ* < 8) are generally more important than higher scales and that scales 4, 1, and 3 are the three most important scales for the classification.

**Fig. 7.**
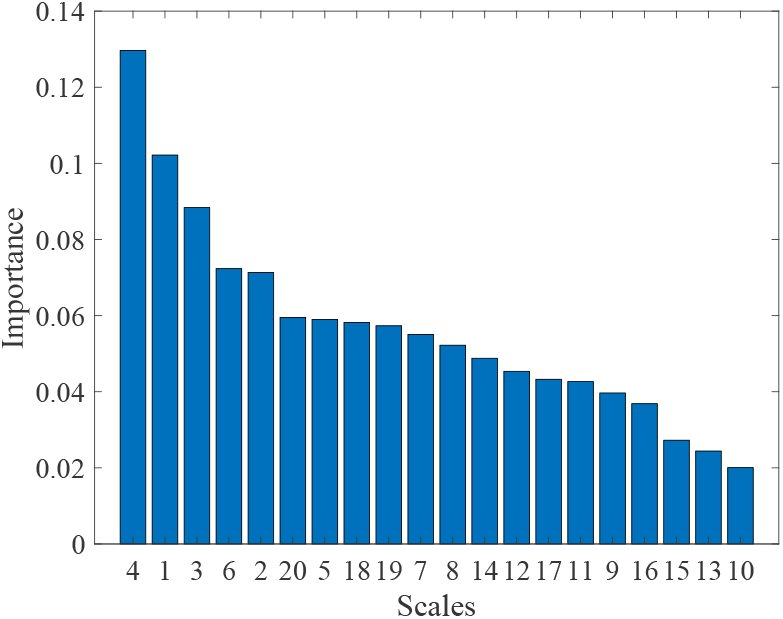
Medians of importance of features over 20 scales.The x-axis is the scale *τ*, and the y-axis is the medians of the normalized importance.

**Fig. 8.**
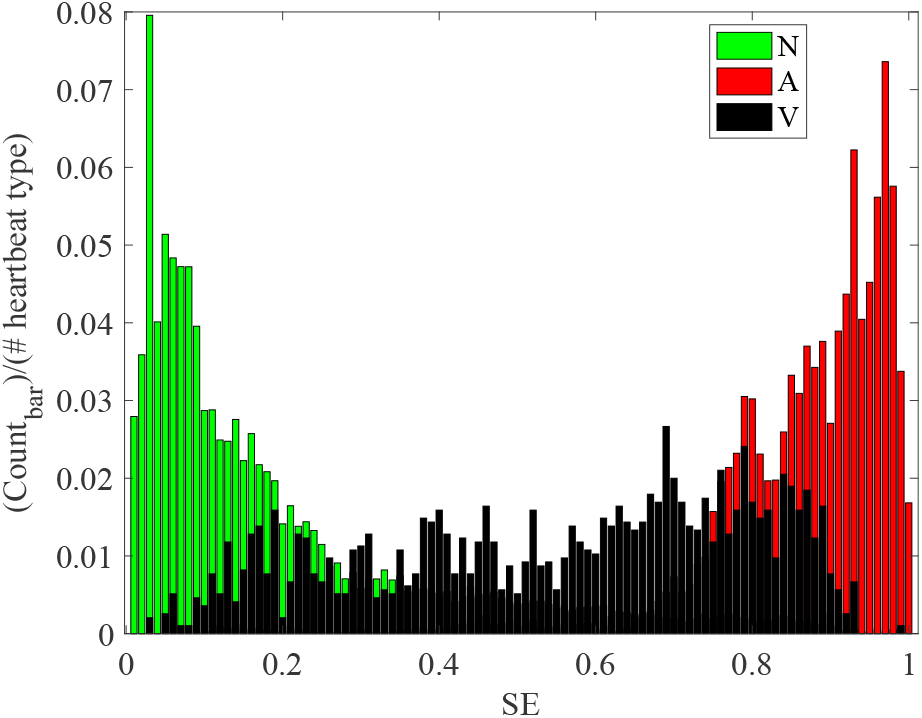
Entropy values of N, A, and V. The x-axis is the Shannon entropy value and, the y-axis is the proportion of the count number of each bar over the total number of each kind of heartbeat.

The Mann-Whitney *U* test was used to test the differences between the importance of two scales for the nine most important scales, whose accumulated importance is 0.63. The results show that the most important scale (4) is significantly different (*p* < 0.05) from all other scales and the second most important scale (1) is significantly different (*p* < 0.05) from the other scales except for scale (3), which is significantly different (*p* < 0.05) from the scales that are less important than itself.

## IV. DISCUSSION

The HbI-based heartbeat classification stands out in the applications based on wearable devices and unconstrained measurements of the cardiac electrical signal, which has poor quality and is prone to be noisy. However, a signal entropy value in the original scale is not adequate in separating different heartbeats. In a follow-up experiment based on the algorithm of Zhou *et al*. [12], we tried to use the algorithm to separate N, AF, and V. We obtained similar results for the binary classification of N and A by setting the threshold of entropy value to 0.63. However, the single entropy value became insufficient when V was added. Fig. 9 shows the distribution of the Shannon entropy values of the three types, from which it can be seen that V sprawls along the x-axis and makes the classification improbable.

There are some hyperparameters to set in our study. A parameter that should be mentioned is the threshold for heartbeat density. *σ_AF_* and *σ_V_* are determined on the basis of the preprocessing of A and V trying to preserve the samples as much as possible. Generally, by lowering *σ_AF_* and *σ_V_*, the sample numbers will increase for AF and V. A natural concern is whether the low densities will cause lower classification outcomes. We thus tried a grid comparison by setting *σ_V_* in [0.10:0.70] with a 0.10 interval and *σ_A_* in [0.05:0.20] with a 0.05 interval, which generated 28 combinations of density thresholds. We then used the combinations to generate AF and V samples which were thrown into the RF model afterward. It is confirmed that little difference can be found with AF and V samples of different thresholds (data not shown here). Noteworthily, greater portions of AF and V samples have density values higher than the ranges we set above. For example, for the 500 HbI, 15% of AF samples are in the range of [0.10:0.70]; whereas the other AF samples have density values higher than 0.70. Similarly, 40% of V samples are in the range of [0.05:0.20]; whereas other V samples have density values higher than 0.20. Therefore, it is reasonable to say the 10% for *σ_AF_* and 5% for *σ_V_* set very low criteria for AF and V and the results of classification suggest that our approach is sensitive to the existence of AF and V.

The underlying idea of this approach is that we extend the target time series to HbI; whereas most studies have focused on the normal sinus heartbeat intervals (NNI) intervals only. That is, all detected R peaks of the ECG signal are included to calculate the beat-to-beat intervals. Therefore, although there may be some ventricular ectopic beats prior to the MAs events, as there are in the SVTDB [29], no special preprocessing is needed to extract the NNI. Moreover, this simplification supports the HbI signal acquired by PPG sensors naturally. This is because, from the PPG signal, normal heartbeats cannot be separated from the abnormal heartbeats that can still pump the arterial blood out to the aorta. By using the HbI instead of NNI combined with the entropy values in multiple resolutions, we can see for the first time the clear differences between the pre-MAs periods and other rhythms.

Intriguingly, despite the MAs and V having different severities, they are similar in the entropy features space, which is consistent with the physiological understanding that the heart experiences ventricular ectopic heartbeats prior to MAs occurring. As for classifying the pre-MAs and V type, we leave this problem to the appropriate machine-learning model. The combination of these two fundamental elements is indispensable for our approach.

We have summarized the previous studies of MAs prediction in TABLE III. Tong *et al.* [22] and Joo *et al.* [24] did not show the precision values of their results. More recently, Joo *et al.* [24], Lee *et al.* [25] and Taye *et al.* [26] used ANN models with ECG signal to construct VT/VF predicting machines. Specially, Getu *et al.* constructed an ANN model with morphological features of ECG signal to predict VT and claimed the accuracy of their approach reached 98.6% [26], however this study only detected VT. Compared with the studies above, our approach predicts two types of MAs (VT/VF) and achieves the highest overall accuracy, sensitivity, and specicity. Moreover, our approach can predict the occurrence of MAs at the earliest, 374 seconds prior to the event.

**TABLE III.** COMPARISON OF PREVIOUS WORKS

| | Required signal | Method | Target | Precision (%) | Recall (%) | Specificity (%) | $time_{max}$ (s) |
| --- | --- | --- | --- | --- | --- | --- | --- |
| Thong et al (2007) [22] | HRV from ECG | Statistical inference based on heart rate | VT<br>VF | - | 53.0 ~ 69.0 | 91.0 | 30 ~ 116 |
| Segyeong Joo et al (2012) [24] | HRV from ECG | ANN model based on HRV metrics | VT<br>VF | 76.6<br>92.2 | 82.9<br>88.9 | 71.4<br>92.9 | 10 |
| Hyojeong Lee et al (2016) [25] | HRV from ECG | ANN model based on HRV metrics | VT | 82.4 | 88.2 | 82.4 | 180 ~ 300 |
| Getu Tadele Taye et al (2019) [26] | HRV from ECG and respiratory rate variability | ANN model based on morphological features of ECG | VT | 98.6 | 98.4 | 99.04 | 30 |
| Proposed approach | HbI | Tree assembly based on complexity of HbI | VT<br>VF | **99.87**<br>**99.87** | **99.99**<br>**99.24** | **99.98**<br>**99.99** | **374** |

The results of the RCMsEn computation and the multi-classification based on the RF model substantiate the underlying assumption about the intrinsic differences among the rhythms of interest. Although the AF and V are the most common types of arrhythmias, a wider spectrum of arrhythmias/cardiac problems should be included. Efforts have been made to include more types of arrhythmias into our model. Specially, we have tried to include atrial flutter (AFL) and AV junctional rhythm (J) from the Atrial Fibrillation Database (AFDB) [30]. However, the AFDB contains only the beginnings of the records with few targeted arrhythmias (AFL: 14 episodes and V: 18 episodes with fewer than 1,000 heartbeats). Despite the limitation due to data unavailability remaining for the time being, we believe that a more comprehensive validation will become possible in the near future.

## V. CONCLUSIONS

In this study, assuming that intrinsic characteristics of the malignant ventricular arrhythmias (MAs) can be reveiled by proper metrics of signal complexity, we proposed an approach combining an improved multi-scale entropy with a nonlinear random forest model to distinguish MAs from other heart rhythms (normal sinus rhythm, atrial fibrillation, and ventricular ectopic contraction) using a relatively short heartbeat intervals. Our approach shows that the heart behaves differently prior to MAs occurring. By using the 500 heartbeat intervals 374 seconds (80 bpm) prior to the MAs, our approach can predict impending MAs accurately (recall=99.99; precision=99.87 for VT; recall=99.24; precision=99.87 for VF). Furthermore, this approach can also detect the atrial fibrillation and ventricular ectopic contraction accurately and is very sensitive to the existence of these arrhythmias. By using this approach, a further study with a wider spectrum of arrhythmias of different pathological origins may be of great value for the systematically understanding the arrhythmias in terms of complexity so as to make wearable/personal heart tracking technologies, e.g., the photoplethysmogram (PPG) sensor, invaluable in heart healthcare.

## ACKNOWLEDGMENTS

This research and development work was supported by the MIC/SCOPE #192207006.

## Footnotes

1 http://physionet.org/physiobank/database/mvtdb/

2 https://physionet.org/content/mitdb/1.0.0/

## Notes

Manuscript received xx, 2019. This work was supported by the MIC/SCOPE 192207006. M. Huang is the Correspondent author of this paper.

